# MOTS-c Coordinates Inter-Organellar Proteostasis for Adaptation to Chronic Metabolic Stress

**DOI:** 10.64898/2026.06.22.733272

**Authors:** Conscience Princesse Bwiza, Ella Schwab, Lily Xia, Ariana Chen, Elijah Song, Sabrina Lin, Erin Kim, Yui Hashiyada, Jyung Mean Son, Michelle Rice, Jessica Kim, Sara Martins, Eun Hee Koh, Berenice Benayoun, Changhan Lee

## Abstract

Mitochondrial communication coordinates adaptive responses across organelles to sustain cellular homeostasis, a network that declines with age and contributes to loss of proteostasis. Here, we identify MOTS-c, an exercise-induced mitochondrial-derived peptide (MDP) encoded within the 12S rRNA locus, as an inter-organellar arm of the mitochondrial stress response (MSR) that links mitochondrial signaling to endoplasmic reticulum (ER) proteostasis and enables adaptation to chronic stress. Using progressive stress media (PSM), a model of gradual and multifactorial metabolic stress, we show that MOTS-c enables adaptation through a biphasic program: acutely, a reversible, ATF4-independent suppression of protein synthesis; and chronically, an ATF6-biased ER unfolded protein response (UPR^ER^) with tempered ATF4 engagement and coordinated metabolic remodeling. Whereas mitochondrial unfolded protein response (UPR^mt^) pathways have been extensively characterized in acute, genetic, and sustained models of mitochondrial perturbation, this work reveals how mitochondrial communication actively engages ER proteostasis during progressive and persistent metabolic stress. By expanding proteostatic capacity while tempering terminal stress signaling, MOTS-c enables cells to withstand chronic stress. Together, these findings define a MOTS-c-dependent arm of the MSR that integrates mitochondrial communication with ER proteostasis to promote chronic metabolic stress adaptation.

## Introduction

Cellular homeostasis is supported by mitochondrial communication that senses metabolic and environmental cues and coordinates adaptive responses across the nucleus, ER, lysosomes, and other organelles through contact sites and signaling pathways (*1-3*). These networks regulate programs beyond bioenergetics, including metabolism, proteostasis, innate immunity, and stress adaptation (*1, 4*). During aging, their fidelity declines, weakening inter-organelle coordination and contributing to mitochondrial dysfunction as a central, interconnected hallmark of aging (*5*).

The mammalian mitochondrial stress response (MSR) is a transcriptional program that induces nuclear-encoded mitochondrial factors in response to mitochondrial perturbation (*1, 6*). One well-characterized arm of this response is the mitochondrial unfolded protein response (UPR^mt^), which preserves mitochondrial proteostasis. Although mitochondrial stress communication is conserved across species, its molecular framework differs. In *C. elegans*, the UPRmt is mediated by ATFS-1, which shifts from mitochondrial import to nuclear accumulation during stress and activates mitochondrial protective genes (*7*). In mammals, mitochondrial stress signaling is more broadly routed through the integrated stress response (ISR), converging on the eIF2α–ATF4 axis (*8-10*). However, mammals also retain ATFS-1-like features of MSR, as mitochondrial depolarization can drive GPS2 translocation from mitochondria to the nucleus, where it activates stress-responsive genes that overlap with ATF4-regulated programs (*11*). Notably, the ATF4-centered MSR/ISR axis is well suited for resolving severe or transient mitochondrial insults, but sustained activation can promote apoptosis rather than durable tolerance (*12*). How mammalian cells preserve proteostasis under chronic, sub-lethal mitochondrial and metabolic stress, a persistent burden likely to accumulate during aging, remains unresolved.

Mitochondrial-derived peptides (MDPs) have emerged as key mitochondrial communicators (*13*). Encoded by short open reading frames in the mitochondrial genome, MDPs act locally and systemically to enhance stress resistance, regulate metabolism, and support organismal resilience (*13*). MOTS-c, a 16-amino-acid microprotein encoded within the mitochondrial 12S rRNA gene, is induced by exercise in skeletal muscle, hypothalamus, and circulation, and, when administered to mice, enhances physical capacity, attenuates age-related muscle loss, improves insulin sensitivity, and improves healthspan (*14-16*). Under metabolic stress, MOTS-c translocates to the nucleus and regulates adaptive nuclear gene expression (*17*), establishing a mitochondria-to-nucleus signaling axis in which the messenger is itself encoded by the mitochondrial genome. *In vivo*, the adaptive effects of MOTS-c are most evident under prolonged, moderate physiological stress, including exercise, a key inducer of endogenous MOTS-c (*14-19*).

Here, we identify MOTS-c as an inter-organellar arm of the mammalian mitochondrial stress response (MSR) that links mitochondrial communication to endoplasmic reticulum (ER) proteostasis, enabling adaptation to the progressive, moderate stress burden associated with aging. We show that MOTS-c adds a distinct adaptive layer to the mammalian MSR, unfolding through temporally separable phases. Acutely, MOTS-c imposes a reversible pause in global translation independently of the canonical ATF4-linked mitochondrial stress pathway. As stress persists, this response transitions toward a tempered convergence with ATF4, favoring adaptation rather than terminal stress signaling. A defining feature of this MOTS-c-induced MSR is strong engagement of the ER, with preferential activation of ATF6, an ER stress-responsive bZIP transcription factor within the broader ATF family that also includes ATF4 and ATF5 (*20*). Using progressive stress media (PSM), an autologous spent-medium model that captures bidirectional, multi-axis metabolic stress rather than acute, unidirectional nutrient withdrawal, we find that MOTS-c coordinates a mitochondrion–nucleus–ER program in which early translational restraint gives way to an ATF6-biased transcriptional and metabolic program marked by significant metabolic remodeling. Together, these findings position MOTS-c as an MSR arm that couples mitochondrial stress signaling to ER proteostasis during prolonged stress, consistent with its role in exercise adaptation, a physiological state that induces MOTS-c expression in humans (*16*).

## Results

### MOTS-c drives adaptation to prolonged metabolic stress

Aging reflects the gradual erosion of cellular homeostasis under persistent, low-grade stress. This burden is progressive and composite, shaped by coordinated changes in nutrient availability, metabolite flux, and intercellular signaling. Yet conventional metabolic stress models often reduce this complexity to a single axis of nutrient loss, such as glucose or serum withdrawal.

To model this complexity, we developed a progressive stress medium (PSM), an autologous spent-medium paradigm in which cells condition their own environment during metabolic stress (**Fig. 1A**). We first defined three conditioning states across a graded stress continuum: a mild state with negligible impact on viability, a severe state associated with terminal decline, and a moderate state that produced addressable stress (**Fig. 1B**). This moderate condition was selected for subsequent studies and designated PSM. Unlike dilution-based nutrient restriction, which simply subtracts nutrients from complete medium, PSM reflects both depletion and accumulation events generated by the cells themselves, creating an integrated metabolic environment shaped by positive and negative metabolite shifts (**Fig. 1C**). In this way, PSM captures a stress window that is biologically consequential but not yet irreversible, remaining initially adaptive while progressively compromising long-term survival (**Fig. S1A–E**).

**Fig. 1.**
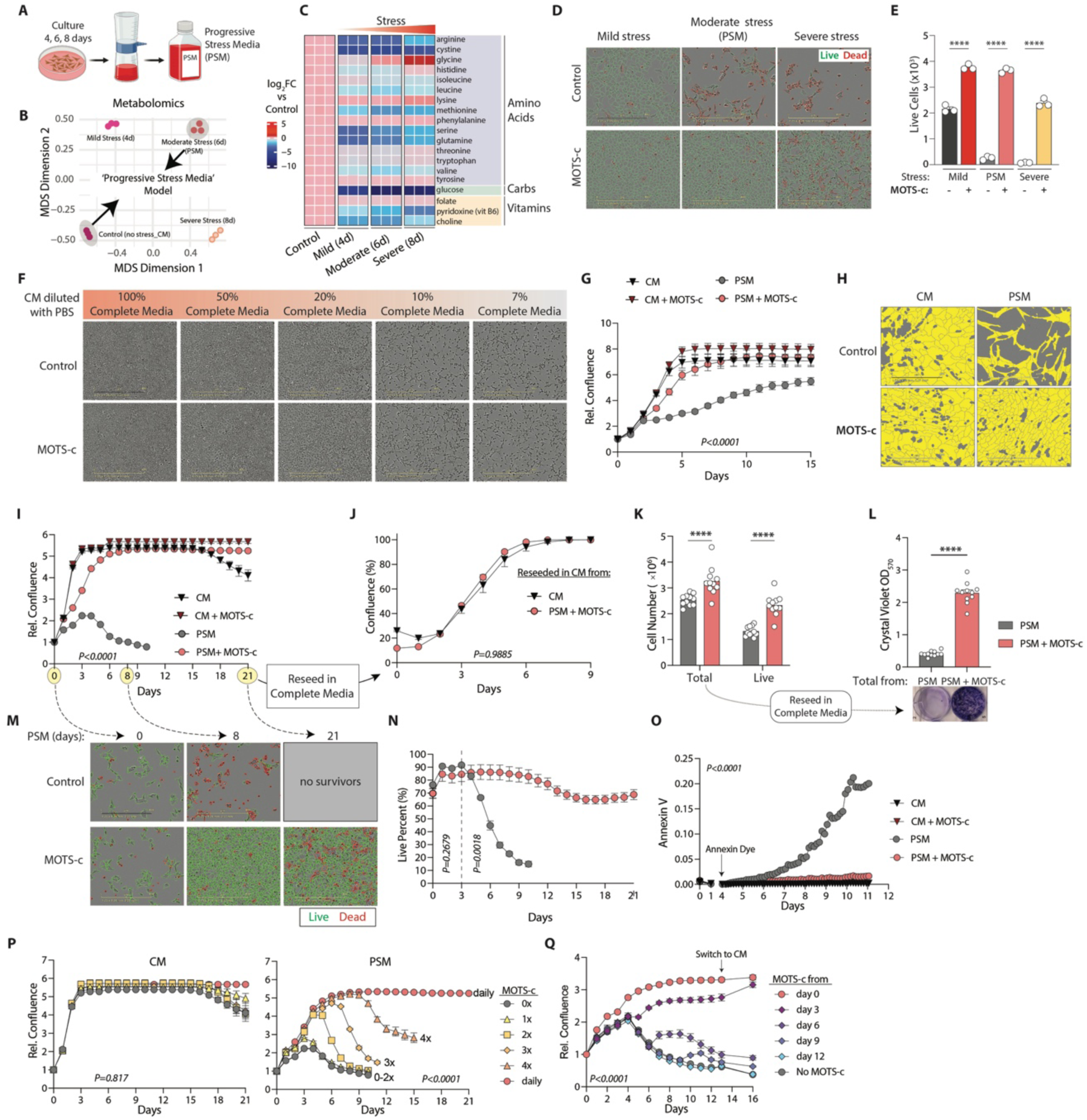
MOTS-c enables chronic adaptation to progressive metabolic stress. (**A**) Schematic of the autologous progressive stress media (PSM) model, in which media collected from HEK293 cultures after 4, 6, or 8 days of growth produces graded, multi-axis nutrient turnover. (**B**) Multidimensional scaling (MDS) of media metabolomes from complete medium (CM), mild stress media, moderate stress media/PSM, and severe stress media. (**C**) Heatmap showing log₂FC versus control for essential media metabolites across stress stages, including amino acids, glucose, folate, vitamin B6, and choline. (**D**) Representative Live/Dead images of HEK293 cells under mild, moderate/PSM, and severe stress, ± MOTS-c on day 6. Green indicates live cells and red indicates dead cells. (**E**) Quantification of live cells from (**D**) across mild, PSM, and severe stress, ± MOTS-c. (**F**) Representative images of cells cultured in diluted CM, 100%, 50%, 20%, 10%, or 7% complete medium in PBS, ± MOTS-c. (**G**) Relative fibroblast confluence over time under CM ± MOTS-c and PSM ± MOTS-c. (**H**) Representative yellow-masked fibroblast images from (**G**) under CM and PSM, ± MOTS-c on day 5. (**I**) Long-window relative confluence over 21 days, with arrow indicating reseeding into CM. (**J**) Confluence of cells reseeded into CM after PSM, CM, or CM + MOTS-c exposure, showing proliferation across groups. (**K**) Total and live cell counts for PSM versus PSM + MOTS-c, with arrow indicating reseeding into CM. (**L**) Crystal-violet OD₅₇₀ of cells reseeded from PSM ± MOTS-c total-cell populations into CM, with representative image of stained wells. (**M**) Representative Live/Dead images of PSM ± MOTS-c cultures from (**I**) on days 0, 8, and 21; day-21 PSM cultures show no surviving cells. (**N**) Live-cell percentage over 21 days, showing collapse of PSM-only cultures beginning around day 9 and rescue by MOTS-c. (**O**) Annexin V apoptosis tracking over 12 days, with annexin dye added on day 4. (**P**) Dose-frequency rescue under CM and PSM, showing relative confluence after 0x, 1x, 2x, 3x, 4x, or daily MOTS-c administration. MOTS-c has no effect under CM but dose-dependently rescues confluence under PSM. (**Q**) Timing-dependent rescue with switch to CM on day 12. MOTS-c was initiated on day 0, 3, 6, 9, 12, or never; full recovery requires MOTS-c delivery before a death checkpoint. Data presented as mean ± SEM. E: Ordinary one-way ANOVA. L: Two-tailed Mann-Whitney test. G, I, J, K, N, O, P, Q: Two-way ANOVA analysis. ****P<0.0001.

MOTS-c significantly improved cellular adaptation and resilience under PSM (**Fig. 1D, E, Fig. S1F**), but not under matched dilution-based nutrient deficiency (**Fig. 1F**), indicating that its activity is not a generic response to nutrient scarcity. We also supplemented PSM with individual nutrients present in complete medium, including glucose, GlutaMax as a glutamine source, and FBS. Among these, glucose produced only a partial rescue and did not restore viability to the level achieved by MOTS-c (**Fig. S1G**). Similar protection was observed in primary human dermal fibroblasts (**Fig. 1G, H**), demonstrating that this response is not restricted to HEK293 cells.

MOTS-c conferred exceptionally prolonged protection under PSM. Using continuous, real-time live-cell imaging to track proliferation and viability, we found that cells cultured in PSM alone proliferated modestly for approximately 3 days before growth ceased (**Fig. 1I**). This arrest was followed by an abrupt loss of viability beginning around day 4 (**Fig. 1N**), consistent with a transition to apoptosis rather than gradual population decline, as further supported by Annexin V measurements (**Fig. 1O**). In contrast, MOTS-c extended cellular expansion to approximately 9 days, at which time cultures reached confluence and remain viable for at least 3 weeks under continuous PSM (**Fig. 1I, M, N, O, Video S1**). Because senescence is also a chronic stress-response state marked by stress resistance and permanent growth arrest, we asked whether MOTS-c had preserved viability by inducing a senescent-like program. When reseeded, MOTS-c–treated cells expanded as robustly as cells maintained in complete medium (**Fig. 1J**), indicating that MOTS-c preserves proliferative capacity rather than imposing permanent arrest (*i.e.* senescence). This adaptive survival phenotype was further supported by trypan blue exclusion and crystal violet staining (**Fig. 1K, L**).

We next asked whether MOTS-c installs a lasting protective state after a single exposure or instead requires continued stimulation during progressive stress. A single treatment did not improve adaptation, whereas repeated administration enhanced survival in a frequency-dependent manner, beginning with two exposures and reaching maximal protection with daily treatment (**Fig. 1P**). We then tested whether MOTS-c could rescue cells after PSM stress was already underway, focusing on day 3, when untreated cultures began to lose viability. Treatment at this transition point produced significant recovery, whereas protection declined with further delay (**Fig. 1Q**). Notably, nutrient replenishment restored full viability only in cells that had received MOTS-c before entering the pre-death state. Thus, MOTS-c supports a sustained adaptive survival program that requires repeated engagement and must be activated before cells cross a death-commitment boundary.

### MOTS-c reprograms metabolism for adaptation to persistent metabolic stress

Cells treated with MOTS-c adapt to and survive persistent PSM (**Fig. 1**), implying metabolic reprogramming suited to scarcity. Because MOTS-c was first identified as an AMPK-activating mitochondrial-encoded microprotein (*14*) that can enter the nucleus under metabolic stress to induce adaptive genes (*17*), we profiled MOTS-c-treated HEK293 cells under PSM at 24 h, the acute-to-chronic transition, and 72 h, early chronic adaptation. Profiles were similar at 24 h but separated by 72 h, when altered metabolites increased 8.7-fold, from 48 to 418 (**Fig. 2A, B**), indicating that MOTS-c-dependent metabolic reprogramming is primarily a chronic-phase response. A limma likelihood-ratio test across the four conditions resolved four metabolite clusters (**Fig. 2C, D**); two tracked MOTS-c under PSM and were enriched for phosphatidylethanolamine (PE), ether/plasmalogen lipids, and bilayer-property terms (**Fig. 2E**). PE rose while ether and plasmalogen lipids fell (**Fig. 2H**). Within PE, diacyl-PE species increased whereas plasmalogen-PE species declined (24 up vs 3 down for diacyl-PE; 1 up vs 7 down for plasmalogen-PE; **Fig. 2O**), a divergence seen only at 72 h, when 35 of 57 PE species were significant compared with none at 24 h (**Fig. 2L**). Pathway-level analysis likewise ranked PE as the most altered sub-pathway, with 17 of 21 species elevated (**Fig. 2F**), while lipid-subclass analysis showed broad declines in glycosphingolipids, acylcarnitines, and plasmalogens, with PE the leading lipid class that rose (**Fig. 2H**). Together, these changes suggest a membrane state favoring remodeling, autophagy, stress resistance, and reduced proliferative expansion (*21-25*).

**Fig. 2.**
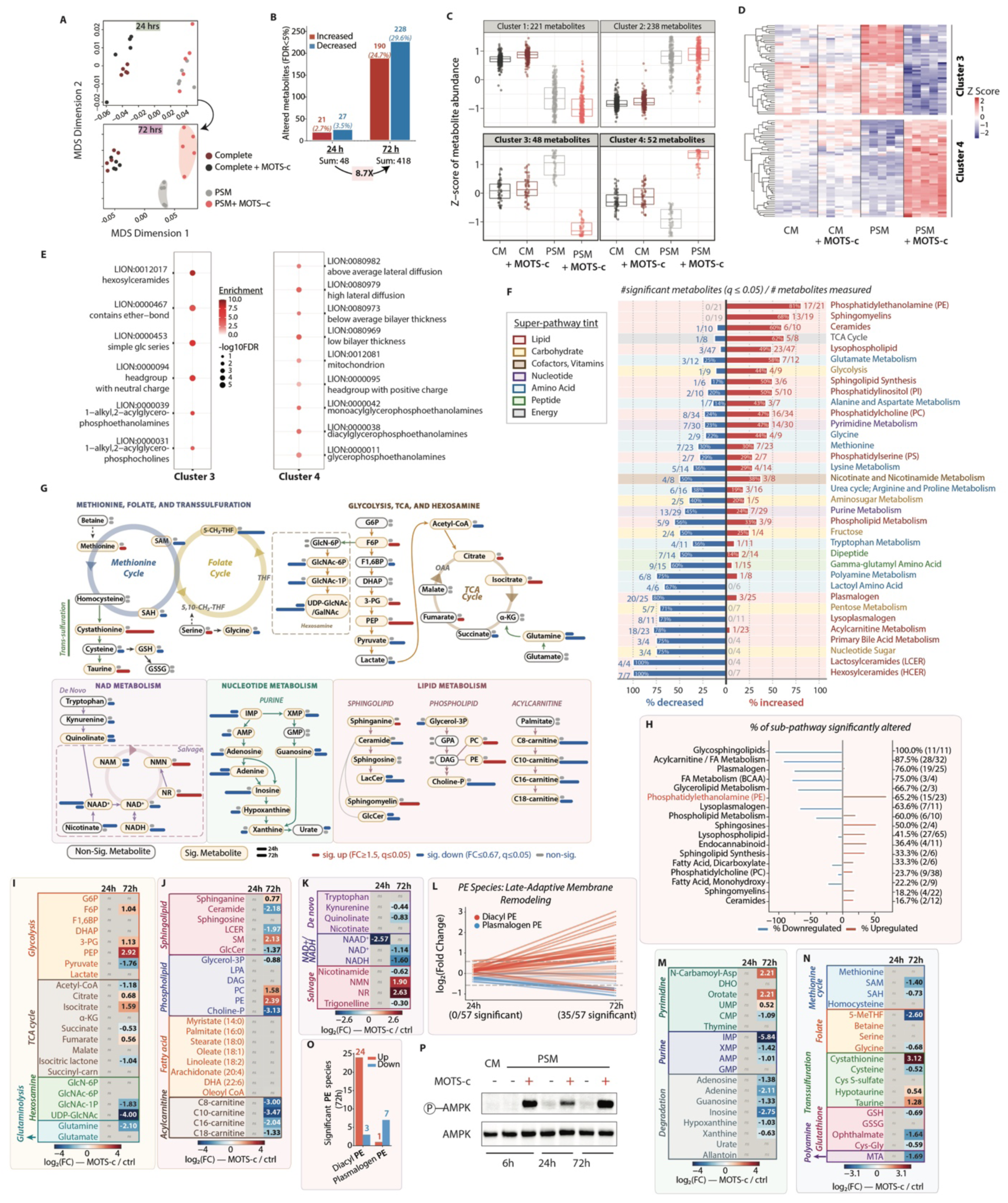
MOTS-c induces comprehensive metabolic reprogramming for persistent metabolic stress adaptation. (**A**) MDS of HEK293 cellular metabolomes at 24 h and 72 h under CM ± MOTS-c and PSM ± MOTS-c, showing predominant late divergence under PSM + MOTS-c. (**B**) Counts of significantly altered metabolites (FDR < 5%) by direction at 24 h and 72 h (48 to 418; 72 h, 228 down / 190 up). (**C**) Cluster-resolved z-score strip plots for four metabolite clusters (Cluster 1: 221; Cluster 2: 238; Cluster 3: 48; Cluster 4: 52 metabolites) across the four conditions. (**D**) Per-cluster heatmap of metabolite abundances (clusters 3 and 4) across CM ± MOTS-c and PSM ± MOTS-c. (**E**) Lipid-ontology (LION) enrichment for Cluster 3 and Cluster 4. (**F**) Sub-pathway ranking of all significant metabolites (q ≤ 0.05) as a fraction of metabolites measured, at 24 h and 72 h, colored by super-pathway tint. (**G**) Integrated metabolic schematic spanning the methionine cycle, folate cycle, transsulfuration, glycolysis, TCA cycle, hexosamine pathway, NAD metabolism, nucleotide metabolism, and lipid metabolism; arrows colored by log₂FC (MOTS-c vs control) at 24 h and 72 h. (**H**) Percent of each lipid subclass significantly altered at 72 h, diverging by direction. (**I**) Glycolysis, TCA cycle, Hexosamine pathway, and glutaminolysis log₂FC heatmap. (**J**) Sphingolipid, phospholipids (PE, PC), fatty acids, and acylcarnitine log₂FC heatmap. (**K**) NAD metabolism log₂FC heatmap. (**L**) Late-adaptive PE remodeling: log₂FC distribution of diacyl-PE vs plasmalogen-PE species at 24 h and 72 h. (**M**) Nucleotide metabolism log₂FC heatmap. (**N**) Methionine cycle and folate/one-carbon, and transsulfuration log₂FC heatmap. (**O**) Counts of significant PE species at 72 h by direction and class. (**P**) Representative p-AMPK Western blot under PSM ± MOTS-c at 6, 24, and 72 h.

Central carbon metabolism followed the same austerity logic. Glycolysis and the TCA cycle coordinate ATP production with redox control (*26*), and by 72 h MOTS-c shifted both toward maintenance. Glycolysis stalled near the phosphoenolpyruvate-to-pyruvate step, with PEP rising 7.6-fold, upstream intermediates accumulating, and pyruvate falling 3.3-fold, consistent with diversion of carbon away from ATP generation (**Fig. 2G, I**) (*27*). Under glucose-limited PSM, glutamine consumption was associated with maintained TCA-cycle citrate pools and redox balance, consistent with a role for glutamine in MOTS-c-dependent anaplerosis, nucleotide metabolism, and sulfur-linked redox buffering (**Fig. 2G, I**) (*28, 29*). Biosynthesis was also restrained: MOTS-c suppressed the hexosamine biosynthetic pathway, with UDP-GlcNAc falling to 0.06-fold (**Fig. 2G, I**), reduced SAM and 5-MeTHF while redirecting methionine flux toward cystathionine and taurine (**Fig. 2G, N**), and contracted total NAD(H) pools while NMN and NR accumulated, indicating reliance on NAD+ salvage over costly de novo synthesis (**Fig. 2G, K**).

Nucleotide and lipid metabolism were similarly curtailed. By 72 h, MOTS-c broadly lowered purine monophosphates, salvage intermediates, and degradation products, indicating reduced purine turnover, while pyrimidine precursors accumulated without downstream nucleotide increases, consistent with an orotate-to-UMP bottleneck (**Fig. 2G, M**). MOTS-c also lowered acetyl-CoA, glycerol-3-phosphate, and acylcarnitines across chain lengths while free long-chain fatty acids remained stable, suggesting restrained mitochondrial fatty-acid import rather than substrate depletion (**Fig. 2G, I, J**) (*30, 31*). Membrane lipid remodeling shifted away from expansion: PE rose more than PC, increasing the PE:PC ratio **(Fig. 2G, H, J)**, while sphinganine and sphingomyelin increased and ceramide, lactosylceramide, glucosylceramide, glycosphingolipids, and plasmalogens declined, consistent with membrane remodeling, autophagy, growth restraint, and altered ether-lipid pools linked to redox stress and ferroptosis sensitivity (**Fig. 2G, J**) (*24, 32*).

Collectively, these coordinated shifts define a MOTS-c-driven austerity program under persistent metabolic stress, integrated with AMPK activation (**Fig. 2P**) (*14, 16, 17*), and optimized for chronic homeostatic adaptation rather than acute growth or simple stress resolution.

### MOTS-c Activates a Transcriptional Program in the Chronic Phase

MOTS-c translocates to the nucleus to orchestrate adaptive transcriptional programs (*17*). To define its transcriptional footprint, we performed RNA-seq under complete medium (CM) and PSM, with and without MOTS-c, at 3h and 24h (**Fig. 3A**). Multidimensional scaling (MDS) showed that MOTS-c-treated cells in PSM followed a distinct transcriptional trajectory that became most apparent by 24 h (**Fig. 3B**), consistent with progressive MOTS-c-dependent reprogramming.

**Fig. 3.**
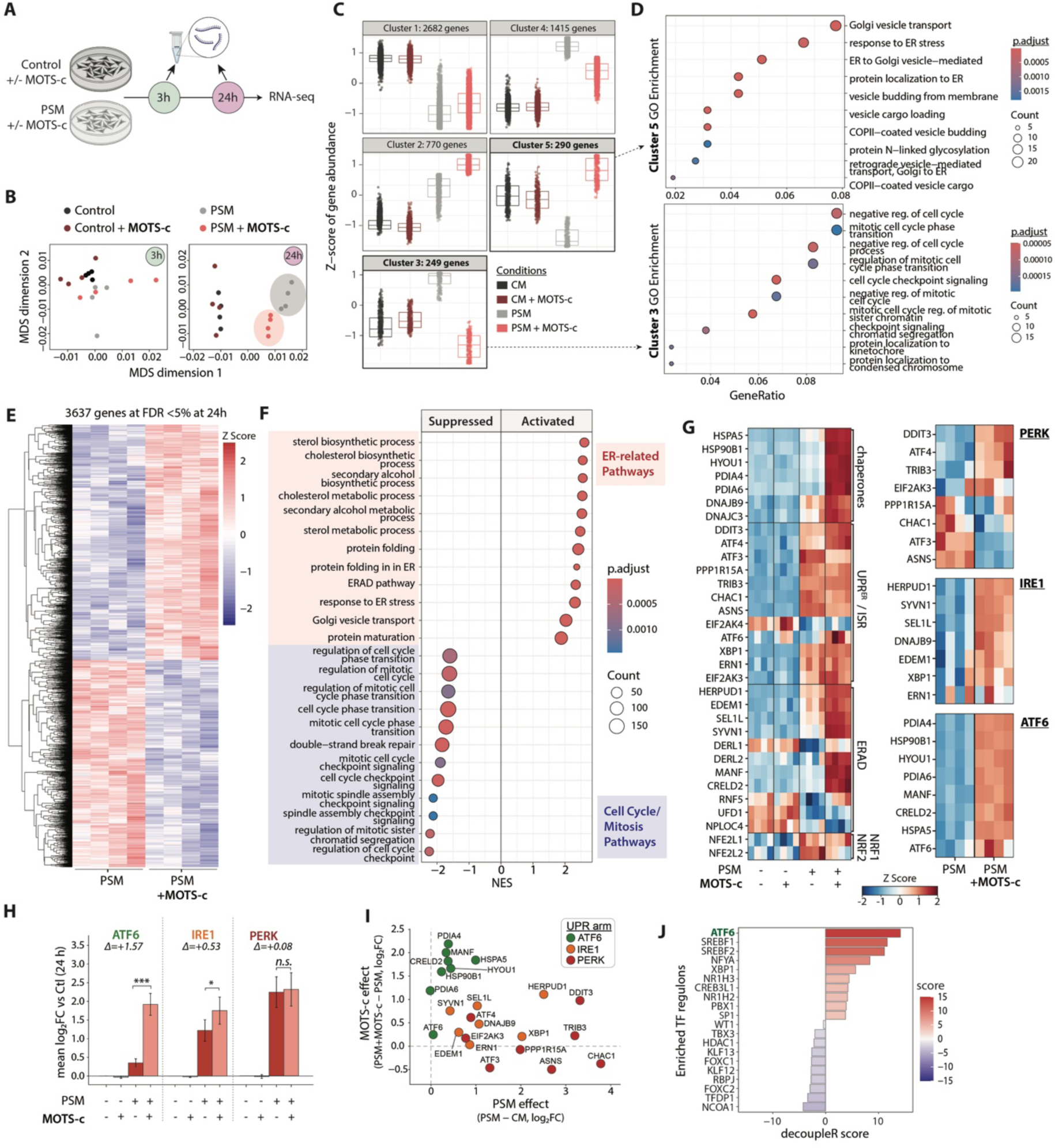
MOTS-c induces a transcriptional program marked by ATF6-biased ER stress response genes. (**A**) Experimental schematic for RNA-seq analysis at 3 h and 24 h under CM ± MOTS-c and PSM ± MOTS-c. (**B**) MDS plot of transcriptomes at 3 h and 24 h, showing progressive divergence of MOTS-c-treated cells under PSM. (**C**) DESeq2 likelihood-ratio test (LRT) cluster-resolved z-score strip plots for five transcriptional clusters at 24 h. (**D**) GO biological process enrichment for Cluster 3, which is repressed under PSM + MOTS-c and enriched for cell-cycle and mitotic-checkpoint programs, and Cluster 5, which is induced under PSM + MOTS-c and enriched for ER stress response, Golgi vesicle transport, and protein N-linked glycosylation. (**E**) Heatmap of 3,637 genes differentially expressed at FDR < 5% in the PSM + MOTS-c versus PSM contrast at 24 h, with all biological replicates shown. (**F**) GSEA dot plot of GO biological processes for the same contrast, separated into activated (ER-related processes; red) and suppressed programs (cell-cycle/mitotic processes; blue). Dot color indicates adjusted p-value and dot size indicates gene-set count. (**G**) Z-scored heatmap of a curated 32-gene proteostasis panel across all replicates and conditions, with branch-resolved heatmaps of canonical PERK, IRE1, and ATF6 target genes. (**H**) Per-arm UPR activation scores were calculated as the mean log₂FC versus control at 24 h for curated ATF6, IRE1, and PERK target genes under PSM or PSM + MOTS-c. The MOTS-c-specific increment was greatest for ATF6 (Δ = +1.57, ***), smaller for IRE1 (Δ = +0.53, *), and minimal for PERK (Δ = +0.08, n.s.). Δ was tested across each arm’s target genes by two-sided paired t-test and confirmed by Wilcoxon signed-rank test. ***P < 0.001, **P < 0.01, *P < 0.05. (**I**) Stress-versus-rescue effect-decomposition scatter of 23 pre-specified UPR effectors. PSM effect (x: PSM − CM, log₂FC) *vs* MOTS-c effect under PSM (y: PSM + MOTS-c − PSM, log₂FC), colored by arm — to test whether MOTS-c boosts effectors not already saturated by stress; PERK/ISR effectors saturate along x while ATF6 chaperones rise along y. (**J**) Unbiased decoupleR analysis of transcription factor regulon activity in PSM + MOTS-c versus PSM at 24 h. Regulons were scored by coordinated changes among their known target genes, without pathway pre-selection.

To identify the programs underlying this shift, we applied DESeq2/LRT. At 3 h, transcripts separated into four clusters primarily defined by medium. By 24 h, five clusters emerged, reflecting both PSM-dependent and MOTS-c-dependent programs (**Fig. 3C**). Two clusters were most prominent under PSM + MOTS-c: Cluster 3, comprising 249 genes, was selectively repressed, whereas Cluster 5, comprising 290 genes, was selectively induced. Gene Ontology enrichment linked Cluster 3 to ER and secretory pathways, including protein folding, ER stress, ER-to-Golgi transport, and COPII vesicle transport, whereas Cluster 5 was enriched for negative regulation of the cell cycle and checkpoint control, including mitotic checkpoint signaling, chromatid segregation, and kinetochore localization (**Fig. 3D**).

Unbiased gene-set enrichment of the PSM + MOTS-c versus PSM contrast at 24 h further identified ER stress and proteostasis as the dominant MOTS-c-induced programs, including protein folding, ER-associated degradation (ERAD), and the response to ER stress. Sterol and cholesterol biosynthesis, which also occur in the ER, were also enriched, whereas cell-cycle and mitotic checkpoint programs were strongly suppressed (**Fig. 3E, F**).

We next asked whether MOTS-c engages ER stress-associated proteostasis programs under PSM, including the canonical UPR^ER^. Using curated gene modules for the PERK, IRE1, and ATF6 arms, along with chaperone, ERAD, ISR, and NRF1/2 programs, we found that PSM + MOTS-c altered these response modules, consistent with broad activation of ER stress and UPR^ER^ pathways (**Fig. 3G**). To determine which branch of the unfolded protein response was most selectively enhanced by MOTS-c, we summarized curated target genes for each UPR arm into an aggregate module score (*33*) and quantified the MOTS-c-specific increase beyond the stress response alone. Mean target expression increased significantly for ATF6 (Δ = +1.57, P < 0.001) and IRE1 (Δ = +0.53, P < 0.05), but not PERK (Δ = +0.08, not significant) (**Fig. 3H**). To separate stress-driven changes from MOTS-c-dependent effects, we plotted each UPR^ER^ gene by its response to PSM alone and its further change with MOTS-c (**Fig. 3I**). PERK/ISR genes were mainly driven by PSM, whereas ATF6-linked ER genes showed stronger additional induction by MOTS-c, indicating an ATF6-biased reinforcement of the UPR^ER^ response. Next, using decoupleR, we inferred transcription factor activity from coordinated changes in known target genes without pre-selecting a pathway. This unbiased analysis identified ATF6 as the most activated regulator in PSM + MOTS-c, ranking above SREBF1/2 and XBP1, further supporting ATF6-linked ER stress remodeling as a major feature of the MOTS-c response (**Fig. 3J**).

Together, these data indicate that MOTS-c initiates a nuclear transcriptional program during the transition from acute to chronic stress, marked by ER proteostasis remodeling and an adaptive, ATF6-biased UPR^ER^ signature.

### MOTS-c Promotes Chronic Stress Adaptation Through Coordinated PERK-ATF6 Signaling

We next explored how MOTS-c affects the three canonical UPR^ER^ arms and their downstream effectors: PERK-ATF4, IRE1-XBP1, and ATF6-BiP. PSM rapidly induced ATF4 within 4 h, whereas MOTS-c suppressed ATF4 induction until 24 h (**Fig. 4A–B**). Notably, CHOP expression increased at 24 h despite delayed ATF4 induction, but returned to comparable levels to PSM by 72 h (**Fig. 4A-B**). MOTS-c may shift the ER stress response from death toward tolerance by raising the threshold for PERK–ATF4 induction while preserving ATF6-mediated proteostasis (*33*), consistent with a model in which the timing and duration of UPR^ER^ branch activity, rather than its mere engagement, dictate whether the response remains adaptive or progresses to apoptosis (*34-36*).

**Fig. 4.**
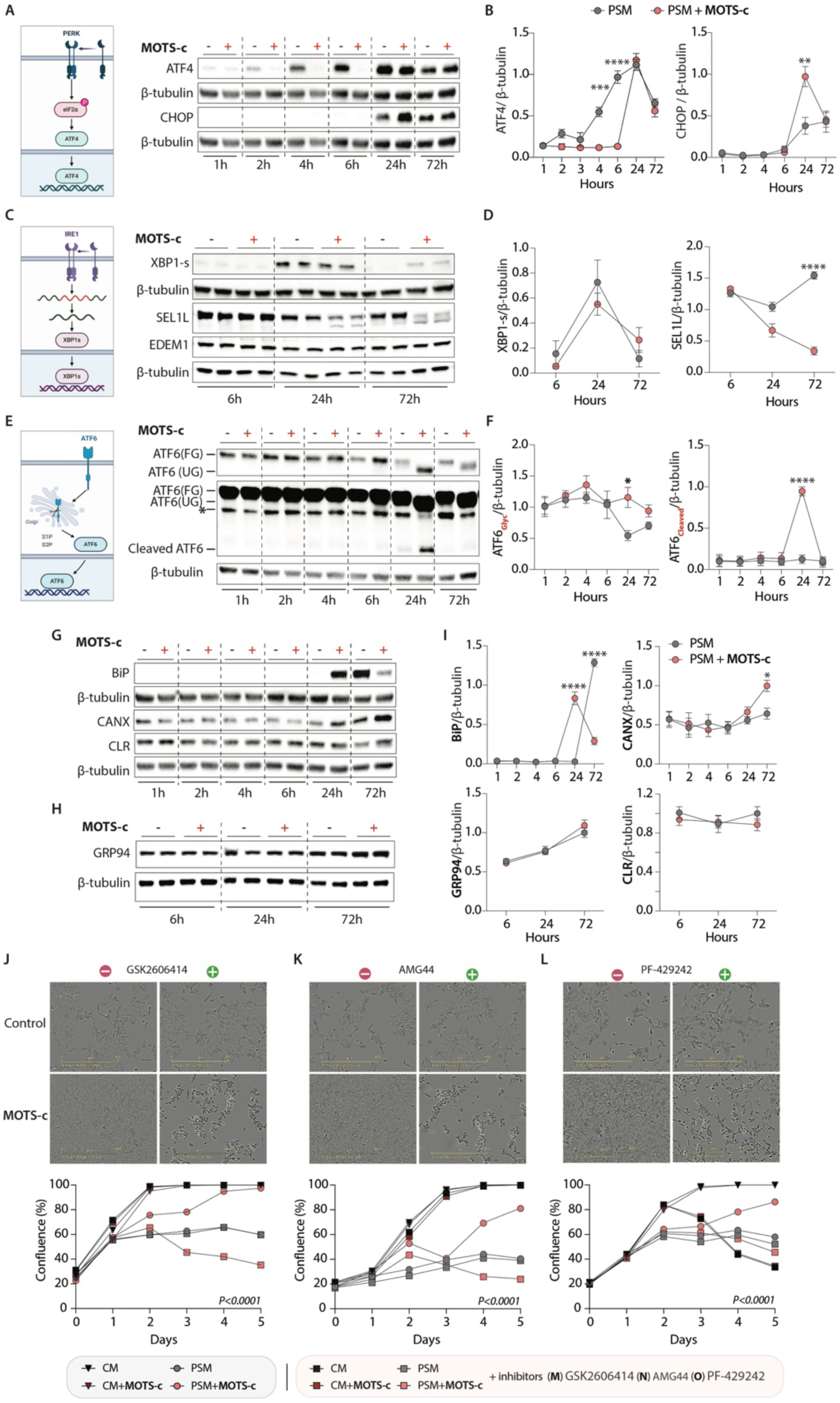
MOTS-c engages the ER unfolded protein response during chronic adaptation to progressive metabolic stress. (**A**) Schematic of the PERK/ATF4 arm and representative ATF4 and CHOP Western blots under PSM ± MOTS-c at 1, 2, 4, 6, 24, and 72 h, with β-tubulin as loading control. (**B**) Densitometric quantification of ATF4/β-tubulin and CHOP/β-tubulin over time under PSM versus PSM + MOTS-c. (**C**) Schematic of the IRE1/XBP1 arm and representative XBP1-s, SEL1L, and EDEM1 Western blots under PSM ± MOTS-c at 6, 24, and 72 h, with β-tubulin as loading control. (**D**) Densitometric quantification of XBP1-s/β-tubulin and SEL1L/β-tubulin over time under PSM versus PSM + MOTS-c. (**E**) Schematic of the ATF6 arm and representative Western blots of fully glycosylated ATF6 P90, cleaved ATF6 P50, and underglycosylated ATF6 under PSM ± MOTS-c at 1, 2, 4, 6, 24, and 72 h, with β-tubulin as loading control. Asterisk denotes non-glycosylated ATF6. (**F**) Densitometric quantification of fully glycosylated ATF6 and underglycosylated ATF6/β-tubulin, and cleaved ATF6/β-tubulin over time under PSM versus PSM + MOTS-c. (**G**) Representative BiP, calnexin, and calreticulin Western blots under PSM ± MOTS-c at 1, 2, 4, 6, 24, and 72 h. (**H**) Representative GRP94 Western blots under PSM ± MOTS-c at 6, 24, and 72 h. (**I**) Densitometric quantification of BiP/β-tubulin, calnexin/β-tubulin, calreticulin/β-tubulin, and GRP94/β-tubulin over time. (**J**) PERK inhibition with GSK2606414 under CM, CM + MOTS-c, PSM, and PSM + MOTS-c, ± GSK. Representative images are shown above, with confluence over 5 days below. GSK2606414 suppresses MOTS-c-dependent protection under PSM. (**K**) PERK inhibition with AMG44, shown as in (J). (**L**) S1P inhibition with PF-429242, which blocks ATF6 cleavage, shown as in (J). Data presented as mean ± SEM and analyzed using two-way ANOVA. K-L: displayed P values corresponding to the effect of treatment condition. ns *P* ≥ 0.123, \**P* < 0.0332, \*\**P* < 0.0021, \*\*\**P* < 0.0002, \*\*\*\**P* < 0.0001.

At the IRE1 branch, MOTS-c did not amplify canonical signaling. XBP1 splicing increased in both conditions, peaked at 24 h, and was modestly lower with MOTS-c before returning toward baseline by 72 h (**Fig. 4C, D**). ERAD machinery showed a distinct pattern. RNA-seq revealed induction of ER quality-control and ERAD genes, including SEL1L and EDEM1 (**Fig 3G**), whereas increased ubiquitination (**Fig. 5C**), reduced SEL1L protein, and sustained EDEM1 abundance suggested post-transcriptional remodeling of the ER proteostasis network.

**Fig. 5.**
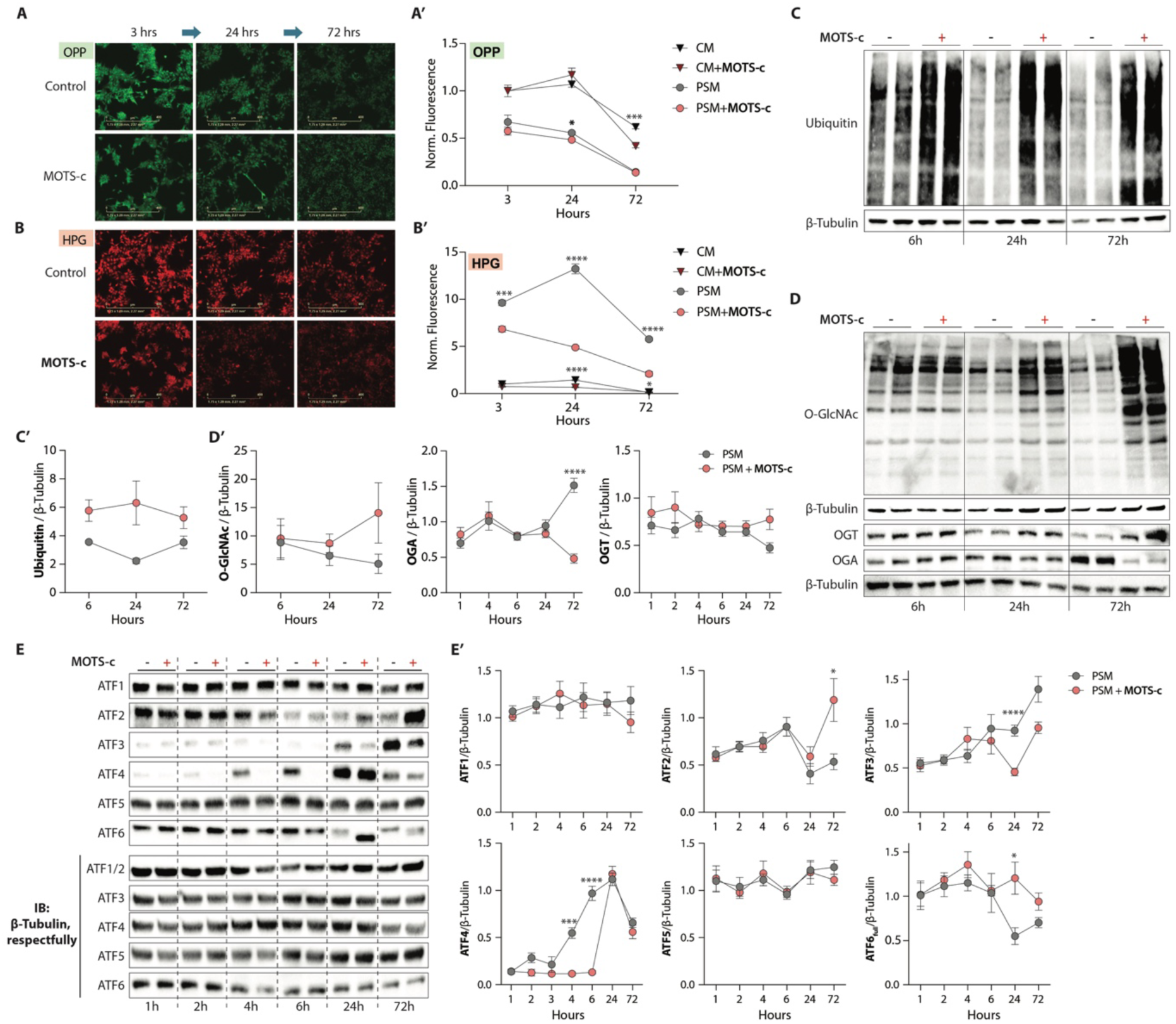
MOTS-c rebalances proteostasis independently of acute ATF4 under progressive metabolic stress. (**A**) Representative O-propargyl puromycin (OPP) immunofluorescence images of nascent translation at 3, 24, and 72 h under control versus MOTS-c conditions. (**A′**) Quantification of normalized OPP fluorescence over 3, 24, and 72 h under CM ± MOTS-c and PSM ± MOTS-c. (**B**) Representative homopropargylglycine (HPG) immunofluorescence images of newly translated proteins at the same timepoints and conditions. (**B′**) Quantification of normalized HPG fluorescence. (**C**) Representative total ubiquitin Western blot under PSM ± MOTS-c at 6, 24, and 72 h, with β-tubulin as loading control. (**C′**) Densitometric quantification of ubiquitin/β-tubulin. (**D**) Representative O-GlcNAc, OGT, and OGA Western blots under PSM ± MOTS-c at 6, 24, and 72 h. (**D′**) Densitometric quantification of O-GlcNAc/β-tubulin, OGA/β-tubulin, and OGT/β-tubulin. MOTS-c preserves global O-GlcNAcylation through increased OGT and reduced OGA despite UDP-GlcNAc depletion. (**E**) Representative ATF1, ATF2, ATF3, ATF4, ATF5, and ATF6 Western blots under PSM ± MOTS-c at 1, 2, 4, 6, 24, and 72 h. (**E′**) Densitometric quantification of ATF1, ATF2, ATF3, ATF4, ATF5, and ATF6/β-tubulin. ATF3 and ATF4 are acutely suppressed by MOTS-c with delayed reappearance in the chronic phase, consistent with ISR redundancy rather than ISR-driven adaptation. Data presented as mean ± SEM and analyzed using two-way ANOVA. ns *P* ≥ 0.123, \**P* < 0.0332, \*\**P* < 0.0021, \*\*\**P* < 0.0002, \*\*\*\**P* < 0.0001.

MOTS-c preferentially engaged ATF6, increasing underglycosylated and cleaved ATF6 (**Fig. 4E, F; Fig. S2A**) and inducing the canonical ATF6 target BiP (**Fig. 4G, I**), an ER chaperone that regulates all three UPR^ER^ sensors. This pattern is consistent with prior work linking ATF6 underglycosylation to ATF6 activation and UPR induction (*37*). The transient CHOP increase at 24 h under MOTS-c may therefore reflect integrated input from multiple UPR^ER^ branches, including ATF6-dependent regulation (*38*). By 72 h, MOTS-c reduced BiP levels (**Fig. 4G, I**), consistent with restoration of proteostatic balance and avoidance of prolonged, apoptosis-prone UPR^ER^ signaling.

Our metabolomics analysis revealed markedly reduced UDP-GlcNAc levels in MOTS-c–treated cells under PSM (**Fig. 2G**). As the key donor substrate for N- and O-glycosylation produced by the nutrient-sensitive hexosamine pathway, reduced UDP-GlcNAc availability is consistent with the increased proportion of underglycosylated ATF6. Defects in N-glycan processing can increase the folding burden on the calnexin/calreticulin cycle; accordingly, MOTS-c treatment elevated ER calnexin levels (**Fig. 4G, I**), but without altering calreticulin (**Fig. 4G, I**) and other chaperones like GRP94 (**Fig. 4H, I**) indicating a selective reinforcement of ER chaperone capacity rather than a broad suppression of the glycosylation machinery.

Functionally, inhibition of ATF6 activation with the S1P inhibitor PF-429242, or inhibition of PERK with GSK2606414 or AMG44, abolished MOTS-c-mediated protection under PSM (**Fig. 4J-L; Fig. S2B**). Indeed, GSK2606414 and AMG44 suppress Tunicamycin-induced ATF4 expression **(Fig. S2C)**. Conversely, PERK activation with CCT020312 restored MOTS-c protection during tunicamycin treatment under PSM (**Fig. S2D**). Together with evidence that PERK–ATF4 signaling can facilitate ATF6 activation and coordinate adaptive UPR responses (*39*), and that ATF6 supports long-term adaptation to chronic ER stress by optimizing ER function (*40, 41*), these data support a model in which MOTS-c promotes stress tolerance through temporally coordinated PERK–ATF6 signaling rather than enhanced IRE1–XBP1/ERAD output.

### MOTS-c sustains proteostasis and imposes a reversible translational pause

Translation is among the most ATP-costly biosynthetic processes and can be attenuated under stress to reallocate energy toward maintenance, consistent with the disposable soma principle (*42*). Using O-propargyl-puromycin (OPP) incorporation, we found that global translation was reduced under PSM and remained low with MOTS-c (**Fig. 5A, A′**). In contrast, methionine-incorporation labeling with HPG in methionine- and cysteine-free medium revealed an additional MOTS-c-specific reduction in nascent synthesis under PSM that was evident by 3h (**Fig. 5B, B′**).

This early effect preceded the chronic phase, occurred before transcriptional and ER-UPR programs were engaged (**Fig. 3**), and was not accompanied by changes in intracellular methionine or SAM, which diverged only by 72 h (**Fig. 2**). Thus, the MOTS-c-dependent decrease in HPG cannot be attributed to downstream metabolic remodeling, but instead, it suggests an acute, primary effect on protein synthesis upstream of later metabolic and ER programs. The low HPG signal in complete medium is expected because replete intracellular methionine competes with HPG incorporation, making the matched PSM comparison the relevant contrast. The selective effect detected by HPG, but not bulk OPP elongation labeling, further suggests that MOTS-c acts at translation initiation or on a subset of nascent proteins rather than broadly suppressing elongation.

This restraint of synthesis was accompanied by a shift toward protein clearance. Total protein ubiquitination increased with MOTS-c relative to PSM alone, reaching significance by 24h and persisting at 72h (**Fig. 5C, C′**). Coupling reduced synthesis with increased ubiquitin conjugation is consistent with active proteostasis rebalancing, in which cells lower biosynthetic and folding demand while marking damaged or misfolded proteins for turnover rather than entering passive synthetic shutdown (*43*). This pattern parallels the transcriptional induction of ERAD and proteasomal programs under PSM + MOTS-c (**Fig. 3F**), suggesting increased engagement of ubiquitin-dependent protein quality control, although direct degradation assays will be needed to determine whether proteasomal flux is increased.

O-GlcNAcylation links nutrient availability to protein quality control by using UDP-GlcNAc to regulate protein stability, ubiquitination, and proteasome function (*44*). UDP-GlcNAc, the end product of the hexosamine biosynthetic pathway and an integrator of glucose, glutamine, acetyl-CoA, and UTP availability, was markedly depleted under PSM + MOTS-c (**Fig. 2G**; 0.06x). Despite this substrate limitation, MOTS-c preserved global O-GlcNAcylation during PSM (**Fig. 5D, D′**), accompanied by increased O-GlcNAc transferase (OGT) and decreased O-GlcNAcase (OGA), the dedicated writer and eraser of this modification, respectively (**Fig. 5D, D′**). Thus, MOTS-c coupled reduced protein synthesis and increased ubiquitin-dependent protein triage with preserved O-GlcNAc signaling, suggesting that proteostatic control is redistributed across complementary post-translational systems even as UDP-GlcNAc availability declines.

Because ATF4 is a central effector of the ISR and an established component of mammalian mitochondrial stress signaling, and because ATF5 has been implicated in the mammalian UPR^mt^, we asked whether MOTS-c-dependent ATF6 induction under PSM was part of a broader ATF response. The ATF family comprises stress responsive bZIP transcription factors that link stress perception to adaptive gene expression. Temporal profiling of ATF1 through ATF6 (**Fig. 5E, E′**) showed that PSM alone most strongly induced ATF4, whereas MOTS-c suppressed this response until 24h. ATF5 changed little. By contrast, ATF6 was again elevated by MOTS-c at 24h and exhibited underglycosylation, consistent with activation. Later changes were also detected in ATF2, which has been associated with inflammatory, cell cycle, DNA damage, and amino acid stress pathways (*45*), and in ATF3, a context-dependent regulator of inflammatory and cell death programs (*46*), whereas ATF1 remained unchanged (**Fig. 5E, E′**). These findings suggest that MOTS-c-dependent ATF6 activation occurs alongside a broader, temporally organized engagement of non-ER ATF factors during chronic stress.

## Conclusion

MOTS-c belongs to a small but growing class of mitochondrial-derived peptides (MDPs), bioactive microproteins encoded within the mitochondrial genome. Encoded within the 12S rRNA, a structural core of the mitoribosome, MOTS-c illustrates that the mitochondrial genome encodes not only components of energy production but also signaling molecules that coordinate stress adaptation. Prior studies have linked MOTS-c to proteostatic and translational control through several contexts, including HSF1-dependent stress adaptation in skeletal muscle (*16*), mTORC1 regulation in immune cells (*18*), and CRIF1-associated mitoribosomal stress signaling in hypothalamic neurons (*15*). Here we extend this framework by showing that an MDP embedded in the mitochondrial translation apparatus (*i.e.* 12S rRNA) regulates cytosolic translation and ER proteostasis, revealing a feedback link between mitochondrial gene expression and cellular protein quality control.

MOTS-c is encoded by the mitochondrial genome, but the adaptive program described here is implemented through nuclear gene regulation (*15-19*). Under persistent mitochondrial and metabolic stress, this program is directed toward the ER, where it strengthens proteostatic capacity. The response proceeds in sequence. MOTS-c first reduces global protein synthesis, lowering the load of nascent proteins entering the cellular folding environment. As stress persists, this early restraint gives way to a nuclear transcriptional program dominated by ATF6-biased ER-UPR signaling, followed by broad metabolic remodeling that supports continued adaptation under PSM. This coupling of early translational restraint to later transcriptional activation resembles the organizing logic of the ISR (*12*), but MOTS-c follows a distinct trajectory: it begins independently of ATF4, later coordinates with ATF4, and sustains ER proteostasis without shifting into the maladaptive signaling associated with prolonged ISR activation and apoptosis (*12*).

This chronic MSR may be particularly relevant to aging (*5*), where stress is cumulative rather than singular. Aging cells experience persistent shifts in metabolism, inflammation, endocrine signaling, and proteostasis that gradually reshape stress sensing and adaptive capacity. We propose that this reflects progressive disruption of mitochondrial stress communication, as coordination among mitochondria, the nucleus, the ER, and other compartments weakens over time. MOTS-c may counter this decline by inducing a prolonged but reversible protective state, promoting adaptation without terminal arrest. These findings position MOTS-c as an inter-organellar mediator of stress resilience and support the mitochondrial endodysbiosis theory of aging proposed here, in which loss of organelle communication drives age-associated homeostatic decline.

## Acknowledgments

This work was supported by the AAUW International Fellowship to C.P.B., the National Institute on Aging (NIA) T32 AG052374 and F31 AG082606, and the Diana Jacobs Kalman/AFAR Scholarship for Research in the Biology of Aging from the American Federation for Aging Research (AFAR) to M.C.R.; the AADOCR Student Research Fellowship to J.S.K.; a Larry L. Hillblom Foundation fellowship to J.M.S.; NIA R01 AG076433, Pew Biomedical Scholar Award #00034120, and the Kathleen Gilmore Biology of Aging Research Award to B.A.B.; and NIA R01 AG052558 and R01 AG084214, NIGMS R01 GM136837, Hevolution, and the Hanson-Thorell Family to C.L.

## Supplemental figures

**Fig. S1.**
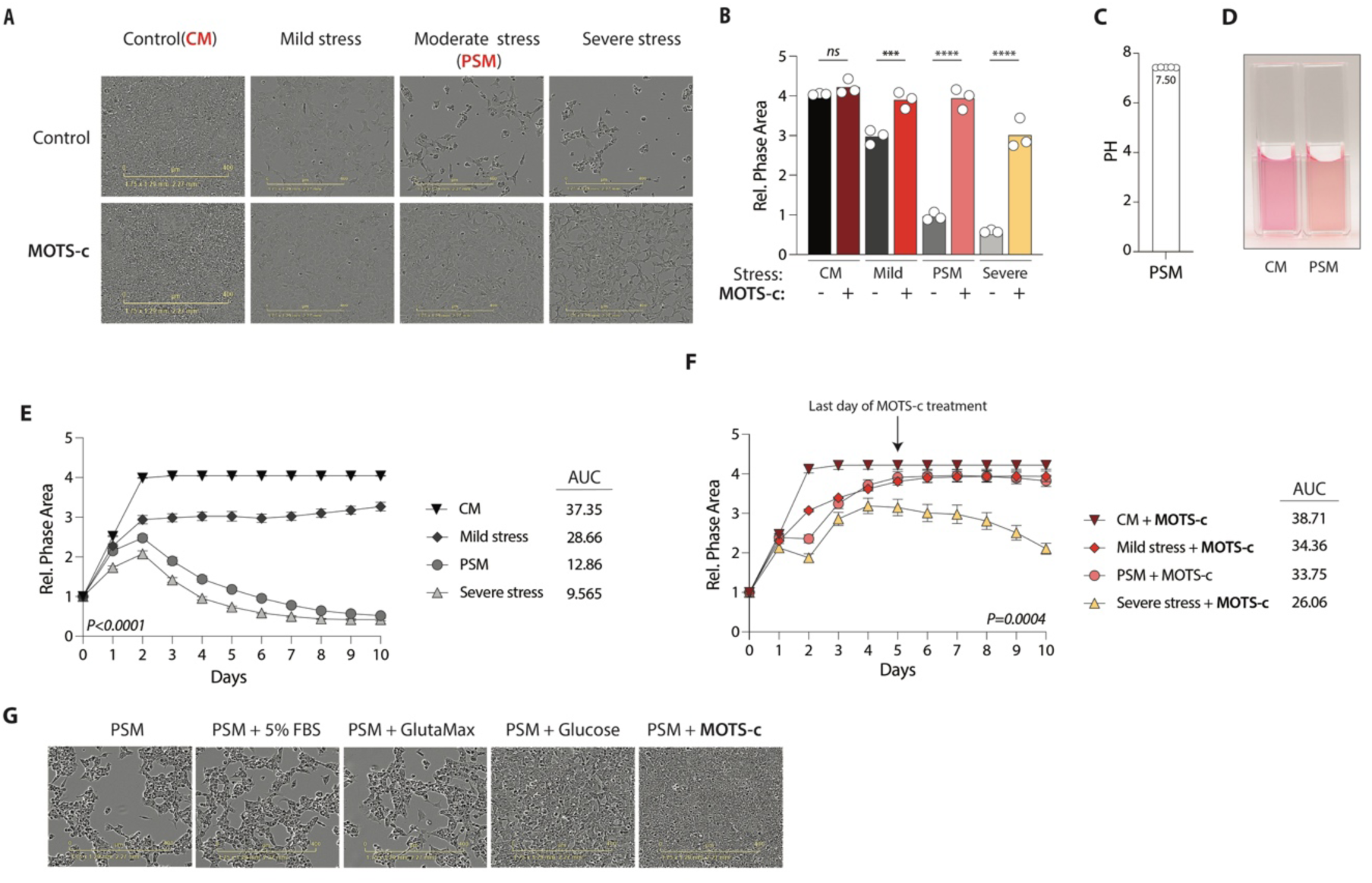
MOTS-c enables chronic adaptation to progressive metabolic stress. (**A**) Phase-contrast images corresponding to Fig. 1D, showing cells under control/CM, mild, moderate/PSM, and severe stress media, each ± MOTS-c. (**B**) Quantification of phase area across CM, mild, PSM, and severe stress ± MOTS-c. MOTS-c significantly increased phase area under PSM, with no significant effect under CM. (**C**) pH of PSM measured on the day of media collection. (**D**) Representative images of CM and PSM shown side-by-side to illustrate the color difference. (**E**) Relative phase area over 10 days under CM, mild, PSM, and severe stress media without MOTS-c. Cells remained viable in mild stress media but not in PSM or severe stress media. AUC is shown inset. (**F**) Relative phase area over 10 days under the same media conditions with daily MOTS-c treatment through day 5, with the final day of MOTS-c treatment indicated. MOTS-c protected cells under mild stress media and PSM, but not under severe stress media. AUC is shown inset. (**G**) Single-nutrient add-back rescue comparing PSM alone with PSM supplemented with 5% FBS, GlutaMax, glucose, or MOTS-c. Glucose partially rescued the PSM phenotype, but not to the extent of MOTS-c. Data presented as mean ± SEM. B: One-way ANOVA. E and F: Two-way ANOVA analysis, with the displayed P values corresponding to the effect of treatment condition. ****P<0.0001.

**Fig. S2.**
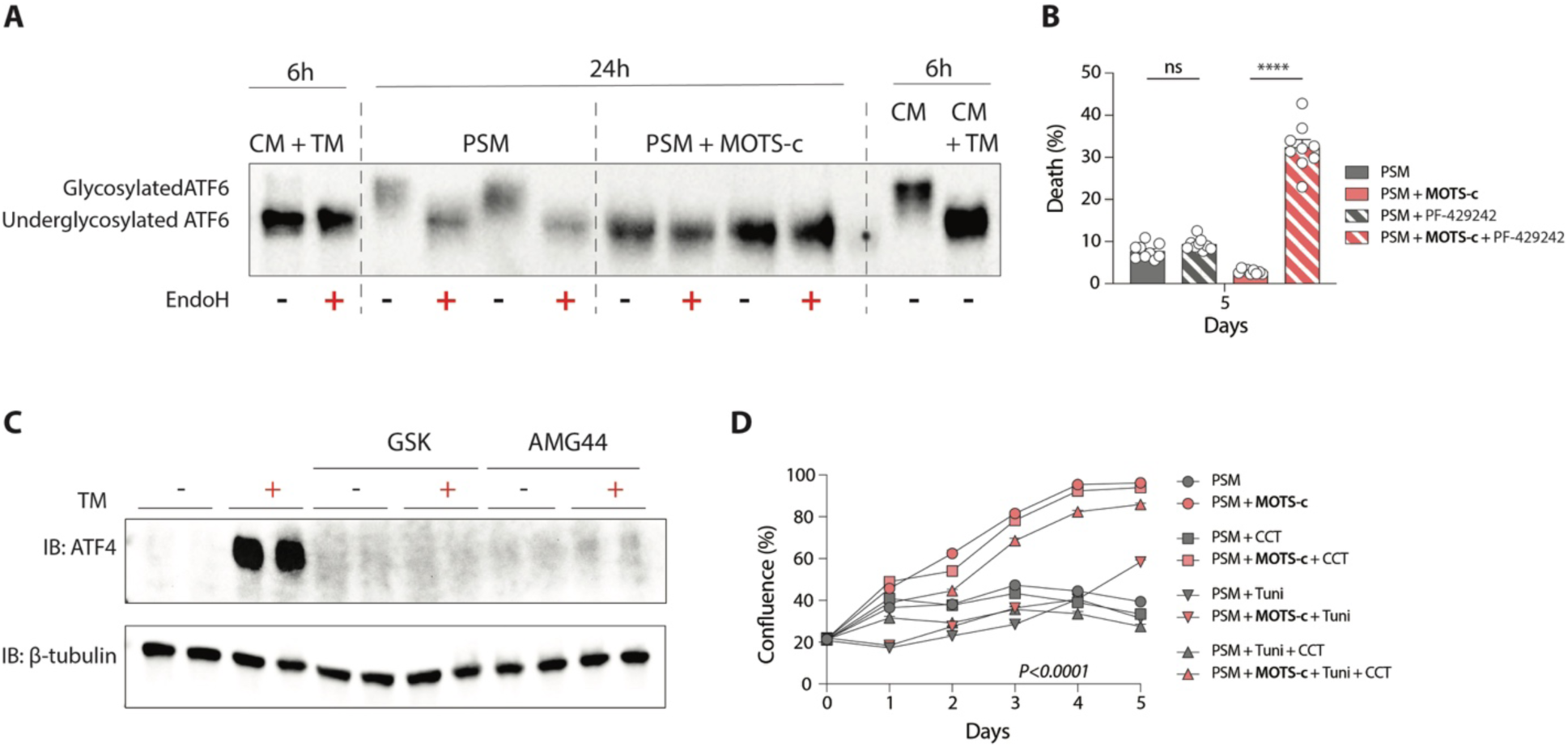
MOTS-c engages the ER unfolded protein response during chronic adaptation to progressive metabolic stress. (**A**) Endoglycosidase H digestion showing that MOTS-c induces ATF6 underglycosylation under PSM, resolving glycosylated and underglycosylated ATF6 at 6 h and 24 h. Tunicamycin served as a positive control for underglycosylation. (**B**) Cell death at day 5 for the conditions shown in Fig. 4L. Death was comparable between PSM and PSM + MOTS-c but markedly increased in PSM + MOTS-c + PF-429242, indicating that S1P/ATF6 inhibition abolishes MOTS-c-mediated protection. (**C**) ATF4 immunoblot showing that the PERK inhibitors GSK2606414 and AMG44 suppress tunicamycin-induced ATF4 expression, confirming on-target PERK inhibition. β-tubulin served as loading control. (**D**) Confluence over 5 days across PSM, ± MOTS-c, ± tunicamycin, and ± the PERK activator CCT020312. PERK activation with CCT020312 restored MOTS-c protection during tunicamycin treatment under PSM. Data presented as mean ± SEM. B: Kruskal-Wallis test; D: Two-way ANOVA analysis, with the displayed P values corresponding to the effect of treatment condition. ns *P* ≥ 0.123, \**P* < 0.0332, \*\**P* < 0.0021, \*\*\**P* < 0.0002, \*\*\*\**P* < 0.0001

## Material and Methods

### Spent Media Production

Cells are seeded at a density of 20×10^6 cells per 10 cm dish in DMEM with 4.5 g/L glucose and glutamine without sodium pyruvate supplemented with 10% FBS (referred to as Complete Media (CM)). The cells are allowed to attach overnight in a cell incubator set at 37°C with 5% CO_2_. The following day, media was replaced, and the cells are cultured to deplete nutrients over periods of 4 days, 6 days, and 8 days. Media is collected at each respective time point, filtered through a 0.45 µm membrane filter, pH recorded (Fisherbrand™ FE150), and stored at 4°C on the day of PSM collection. Media collected on the 4^th^, 6^th^, and 8^th^ days are designated as Mild-stress media, Moderate-stress media (PSM), and Severe-stress media, respectively.

### Proliferation and Survival Experiments

Cells are seeded at a density of 0.1×10^6^ cells and 1×10^6^ per well in 12-well plates and 1×10^4^ cells and 5×10^4^ cells per well in 96-well plates in DMEM containing 4.5 g/L glucose and glutamine, without sodium pyruvate, and supplemented with 10% FBS (referred to as Complete Media (CM)). The cells are allowed to attach overnight in a cell incubator set at 37°C with 5% CO_2_.

The following day (Day 0), we image the cells, and then replace the media with one of the following conditions: CM + H_2_O, CM + 10 µM MOTS-c, Spent Media + H_2_O, or Spent Media + 10 µM MOTS-c. The MOTS-c stock concentration is always prepared at 1 mM in water. We continue daily treatment until the last day of the experiment, depending on the end goal.

Cell proliferation and survival were monitored using the Incucyte® SX5 Live-Cell Analysis System (Sartorius). Phase-contrast images were acquired daily, and cell confluence was quantified automatically using the Basic Analyzer module of the IncuCyte software, which measures the percentage of the imaged area occupied by cells. In addition, cell viability was assessed using the AI Cell Health Analysis, which employs AI-driven cell segmentation and label-free Live/Dead classification to quantify cell health. Raw and normalized confluence and cell health data were exported from IncuCyte and analyzed in GraphPad Prism 10 for data visualization and statistical analysis.

To confirm the viability of the surviving cells at the end of the experiment, the cells are trypsinized and then reseeded in 6-well plates at a density of 5×10^4 total cells per well or 1/5 of cells were reseeded per well in 12-well plates.

For the Trypan Blue exclusion assay, 15 µL of Trypan Blue was mixed with 15 µL of the cell suspension in a tube and gently flicked to ensure thorough mixing. A 15 µL aliquot of the mixture was then transferred onto a counting slide, which was inserted into the Countess instrument to determine the cell concentration (cells/mL).

For the crystal violet assay, cells were incubated with crystal violet for 20 minutes at room temperature on a bench rocker. The stain was then removed, and the cells were washed with tap water until the plate background was clear. Plates were left open at room temperature to air-dry overnight. The following day, an equal volume of methanol was added to each well, and the plate was incubated with its lid on for 20 minutes at room temperature on a bench rocker. The optical density of each well was then measured at 570 nm (OD_570_) using a plate reader.

### Annexin V Assay

Cells were seeded in 96-well plates and after overnight incubation, the CM seeding media was replaced by treatment media ± MOTS-c. 4 days post daily MOTS-c treatment, a fluorescently labeled Annexin V reagent was added to the final concentration of 10 µM and continued daily imaging and MOTS-c treatment. Phase-contrast and fluorescence images were acquired at regular intervals over the indicated time course. Apoptotic cells were identified by Annexin V–associated fluorescence, and signal intensity or positive cell counts were quantified using Incucyte analysis software to assess apoptosis kinetics in real time.

### Western Blotting

Cellular lysates were prepared using RIPA buffer (Thermofisher, Cat#: 89901) containing a protease/phosphatase inhibitor cocktail to prevent protein degradation. The lysates were subsequently sonicated to ensure complete cell lysis and homogenization. The supernatants were subjected to electrophoresis using 8%–16% or 7.5% pre-cast SDS-PAGE gels (Bio-Rad) to resolve the proteins by molecular weight. The resolved proteins were then transferred from the gels to PVDF membranes, blocked using either 5% bovine serum albumin (BSA) or 5% non-fat dry milk in Tris-buffered saline containing 0.05% Tween-20 (TBS-T), and incubated with the appropriate primary antibodies at 4°C overnight. After primary antibody incubation, the membranes were washed three times with TBS-T to remove any unbound antibodies, and then incubated with HRP-conjugated secondary antibodies at room temperature for 1 hour. The membranes were washed three additional times with TBS-T, developed using Amersham ECL Prime Western Blotting Detection Reagent (VWR) or Clarity Western ECL substrates (Bio-Rad), and imaged using the ChemiDoc XRS+ system (Bio-Rad). The bands intensities were quantified using ImageJ.

### Protein Synthesis Assays

The protein synthesis assays were performed using the Click-iT™ Plus OPP Alexa Fluor™ 488 Protein Synthesis Assay Kit (ThermoFisher, Cat# C10456), and Click-iT™ HPG Alexa Fluor™ 594 Protein Synthesis Assay Kit (ThermoFisher, Cat#: C10429) with specific modifications: steps requiring double washes were replaced with single washes using double the volume of the washing solution; following step 5.6, cells were washed once with PBS and then directly proceeded to the Imaging and Analysis step, omitting the DNA Staining step; and Poly-D-Lysine coated plates were used to enhance cell adherence, as we noticed that cells were detaching during the assay process with standard plates. Cell imaging and fluorescence quantifications were done using IncuCyte Orange channel for the Click-iT™ HPG Alexa Fluor™ 594 Kit and Green channel for the OPP Alexa Fluor™ 488 Kit.

### RNA-Seq

Cells were seeded at 1 × 10^5^ cells per well in 12-well plates in DMEM containing 4.5 g/L glucose and glutamine without sodium pyruvate, supplemented with 10% FBS, referred to as complete medium (CM). Cells were allowed to attach for 48 h at 37°C and 5% CO_2_, after which medium was replaced with CM or PSM ± MOTS-c. RNA was extracted after 3 h and 24 h of treatment using the Direct-zol RNA MiniPrep kit.

Paired-end FASTQ reads were pseudoaligned to human GRCh38.p13 transcripts from Ensembl 99 using kallisto 0.43.0 (*47*). Transcript pseudocounts were parsed with a custom Perl script to generate a transcript count matrix for import into R 4.5.0. Transcript counts were summarized to genes, rounded to integers, and analyzed with DESeq2 1.48.0 (*48*). The 3 h and 24 h timepoints were analyzed separately to assess time-dependent effects of MOTS-c. For each timepoint, likelihood-ratio testing (LRT) was performed with each medium and treatment combination modeled as a distinct group. Given the sensitivity of the LRT, significant genes were defined by adjusted p < 10^-3^. Differentially expressed gene clusters were identified using degPatterns from DEGreport 1.44.0. Gene Ontology overrepresentation analysis was performed on significant LRT gene clusters using org.Hs.eg.db 3.21.0 and clusterProfiler 4.16.0, with all DESeq2-detectable genes used as the enrichment background. Terms with false-discovery rate (FDR) < 5% were considered significant. For transcription factor activity prediction, a pairwise DESeq2 Wald-test analysis was also performed at 24 h to estimate the effect of MOTS-c in PSM.

The UPR was divided into its three canonical arms based on published UPR frameworks (*49, 50*). Arm-specific gene sets were curated from orthogonal studies defining branch-selective transcriptional outputs. The ATF6/ER-chaperone set included ATF6, HSPA5/BiP, HSP90B1/GRP94, HYOU1, PDIA4, PDIA6, MANF, and CRELD2; the IRE1α/XBP1s set included ERN1, XBP1, HERPUD1, EDEM1, DNAJB9, SEL1L, and SYVN1 (*51-53*); and the PERK/ATF4 integrated stress response set included EIF2AK3, ATF4, DDIT3/CHOP, ATF3, PPP1R15A/GADD34, TRIB3, CHAC1, and ASNS (*34, 49*). The ATF6-selective assignment was further supported by prior branch-specific analyses (*54*).

Curated UPR/proteostasis heatmaps were generated from DESeq2 LRT variance-stabilizing transformed (VST) counts. The 32-gene proteostasis panel was grouped into ER chaperones, UPR^ER^/ISR effectors and core arm regulators, ERAD machinery, and NRF1/NRF2 modules. Each biological replicate was shown as an individual column and arranged by the 2 × 2 design of PSM and MOTS-c treatment. For each gene, VST values across all replicates were converted to row Z-scores and displayed on a symmetric diverging scale clipped at ±2. Genes were ordered within fixed functional categories without clustering, providing a descriptive view of condition-level shifts across chaperone, ISR, ERAD, and NRF modules. UPR effector cascade heatmaps resolved the curated UPR gene set by canonical arm as three stacked heatmaps: PERK, n = 8 genes; IRE1, n = 7; and ATF6, n = 8. Genes were shown in rows, with the four PSM and four PSM+MOTS-c biological replicates displayed as adjacent column groups separated by a divider. Expression values were taken from VST-normalized counts generated from the same LRT model used to define the differentially expressed gene set.

Saturation scatter plots were generated for 23 pre-specified adaptive UPR effectors comprising the ATF6/ER-chaperone, IRE1, and PERK gene sets. Using the same DESeq2 model, two Wald-model log2 fold-change contrasts were extracted to separate the stress response from the MOTS-c effect under stress. The x-axis represents the PSM effect relative to untreated complete-medium control, and the y-axis represents the MOTS-c boost on the stress background, calculated as PSM+MOTS-c versus PSM. The PSM + MOTS-c versus PSM contrast was the primary MOTS-c-in-stress comparison, while the PSM versus control contrast was extracted from the same model to place both axes on a common statistical footing. P values were adjusted genome-wide using the Benjamini–Hochberg FDR procedure. Each point represents one gene and was colored by UPR arm, with reference lines at x = 0 and y = 0 and non-overlapping gene labels. This two-axis visualization distinguishes genes induced primarily by stress from those further reinforced by MOTS-c under stress.

For transcription factor activity prediction, pairwise DESeq2 results were used as input for decoupleR 2.14.0. TF activity shifts were quantified using the wmean metric, and transcription factors were considered differentially active in the presence of MOTS-c at p < 0.05. The top 20 most significant transcription factors were reported.

### Metabolomics/lipidomics

Cells were seeded at 5 × 10^6 cells per 10 cm dish in DMEM containing 4.5 g/L glucose and glutamine without sodium pyruvate, supplemented with 10% FBS, referred to as complete medium (CM). Cells were allowed to attach overnight at 37°C and 5% CO2, after which medium was replaced with CM or PSM ± MOTS-c. Cells were pelleted and frozen after 24 h or 72 h of treatment, with MOTS-c replenished daily for the 72 h condition. Frozen cell pellets, together with frozen CM, low-stress, PSM, and high-stress media samples, were submitted to Metabolon for untargeted metabolomic profiling.

Metabolon output quantification tables were imported into R 4.5.0 using readxl 1.4.5, yielding 953 metabolites. To address missingness, metabolites with >10% missing values were removed, excluding 131 metabolites, and remaining missing values were imputed using the cart method in MICE 3.19.0 (*55*). Intensities were normalized to PARAM_BRADFORD_MG/ML to correct for sample input and then variance-stabilizing normalized using normalizeVSN from limma 3.64.0, as described previously (*56*). The 24 h and 72 h timepoints were analyzed separately to assess time-dependent effects of MOTS-c. For each timepoint, one-way ANOVA was performed with each medium and treatment combination modeled as a distinct group, and raw p values were adjusted by the Benjamini–Hochberg method using p.adjust. Significant metabolites were defined by adjusted p < 10^-3^. Differentially abundant metabolite clusters were identified using degPatterns from DEGreport 1.44.0, following a strategy similar to prior work (*57*). Metabolon-provided metadata were used to define all QC lipids as the universe/background and to extract lipids within each degPatterns cluster for Lipid Ontology (LION) enrichment analysis (*58*). Enrichments were computed for each cluster using the LION web interface, version 2023.04.14, and terms with FDR < 5% were considered significant.

To report intuitive linear fold changes, Bradford-normalized intensities were log2-transformed and analyzed with limma. The MOTS-c effect under metabolic stress was estimated at each timepoint as the PSM + MOTS-c versus PSM contrast using a moderated t-test, with Benjamini–Hochberg FDR correction across all features. Fold change was calculated as 2^(log2 coefficient), and metabolites were considered significantly changed at FDR ≤ 0.05 with fold change ≥ 1.5 or ≤ 0.67. Complex lipid classes, including ceramides, sphingomyelins, and phosphatidylcholines, were quantified as the summed normalized intensity of all detected species within the corresponding Metabolon sub-pathway. For each metabolite, 24 h and 72 h fold changes were displayed as paired bars on the pathway map, colored by direction of change and scaled to |log2 fold change|. Analyses were performed in R 4.5.3 using limma 3.66.0.

For curated metabolomic pathway heat-strip panels, the MOTS-c effect under metabolic stress was evaluated at each timepoint as PSM + MOTS-c versus PSM + vehicle. For each metabolite, linear fold change was calculated from Bradford-normalized intensities as FC = mean[PSM + MOTS-c] / mean[PSM + vehicle]. Significance was tested on log2-transformed intensities using Welch’s two-sample t-test with unequal variances; metabolites with fewer than two non-missing values per group were excluded. P values were corrected across all detected features within each timepoint using the Benjamini–Hochberg FDR procedure, and fold changes were reported only for metabolites with q < 0.05. Metabolites were grouped into canonical pathways using Metabolon CHEM_ID annotations, including glycolysis, TCA cycle, hexosamine biosynthesis, glutaminolysis, sphingolipids, phospholipids, fatty acids, acylcarnitines, NAD^+^ synthesis/salvage, nucleotide metabolism, one-carbon metabolism, transsulfuration, glutathione, and polyamine metabolism. Complex lipid classes were represented by a single annotated representative species, with paired 24 h and 72 h heat strips shown side by side for each metabolite. Heat-strip analyses and figure rendering were performed in R 4.5.3 using readxl and base R grid.

